# New intranuclear symbiotic bacteria from macronucleus of *Paramecium putrinum* — *Candidatus* Gortzia yakutica

**DOI:** 10.1101/2020.01.13.895557

**Authors:** Alexandra Beliavskaia, Maria Logacheva, Sofya Garushyants, Jun Gong, Songbao Zou, Mikhail Gelfand, Maria Rautian

**Affiliations:** Laboratory of Protistology and Experimental Zoology, Saint Petersburg State University, Russia; Skolkovo Institute of Science and Technology, Moscow, Russia; School of Marine Sciences, Sun Yat-Sen University, Zhuhai, China; Yantai Institute of Coastal Zone Research, Chinese Academy of Sciences, Yantai, China; Faculty of Computer Science, Higher School of Economics, Moscow, Russia; Kharkevitch Institute for Information Transmission Problems, Moscow, Russia

## Abstract

*Holospora*-like bacteria are obligate intracellular *Alphaproteobacteria*, inhabiting nuclei of *Paramecium* ciliates and other protists. *Alphaproteobacteria* have drawn significant attention, as both closest existing relatives of bacteria that gave rise to mitochondria, as well as a class of intracellular bacteria with numerous important pathogens.

HLB clade includes two genera – *Holospora* (Hafkine 1980) and *candidatus* Gortzia (Boscaro 2013). These bacteria have a peculiar life cycle with two morphological forms, a strict specificity to the host species and the type of nucleus they inhabit.

Here we describe a new species of HLB – *candidatus* Gortzia yakutica sp. nov., a symbiont from macronucleus of *Paramecium putrinum*, the first known HLB for this *Paramecium* species. The new symbiont shows morphological similarities with other HLB. The phylogenetic analysis of SSU rDNA gene places it into *candidatus* Gortzia clade.

## Introduction

*Paramecium* ciliates (*Oligohymenophorea*, *Ciliophora*, *Alveolata*) host diverse intracellular symbionts (S. Fokin and Gortz 2009; Preer, Preer, and Jurand 1974; Gortz 2006; Gong et al. 2014), among which the best studied are *Holospora*, obligately intranuclear bacteria of order *Holosporales*, class *Alphaproteobacteria.* Nine *Holospora* species have been morphologically characterized and named so far; however, the 16S rRNA genes have been sequenced for only four of them (*H. obtusa, H. undulata, H. acuminata,* and *H. curviuscula*). *Holospora* have a complex life cycle involving two morphological stages, infectious and reproductive. The infectious forms (IFs) are usually large (up to 20 μm long), have hypertrophied periplasm occupying about one half of the cell, and a recognition tip on the periplasm end. They can survive in ambient conditions for several hours and infect new host cells. Reproductive forms (RFs) are small and able to reproduce by binary fission, and can transform into IFs. These features are used to assign bacteria to the group of *Holospora*-like bacteria (HLB) even when the full sequence of the 16S rRNA gene is not available.

*Holospora* species are divided into two groups according to the behavior of the infectious forms under division of the infected nucleus. Only infectious forms of *H. obtusa*, *H. undulata*, *H. elegans*, *H. curviuscula*, and *H. acuminata* gather near the center of the spindle apparatus of the dividing host nucleus forming the so-called connecting piece, while the reproductive forms mainly appear in the apical parts of the nucleus. Following formation of the connecting piece, IFs escape into the cytoplasm and then into the ambient environment. The second group of *Holospora* species (*H. caryophila, H. bacillata,* and *H. curvata*) does not form connecting piece (Garrity 2000). Boscaro et al. recently have reported a new genus of HLB, *Candidatus* Gortzia, which is evidently a sister taxon to the genus *Holospora* (Boscaro et al. 2013). The genus currently includes two species, *Сandidatus* Gortzia infectiva (Boscaro et al. 2013), and *Сandidatus* G. shahrazadis (Serra et al. 2016), macronuclear symbionts of *P. jenningsi* and *P. multimicronucleatum,* respectively. The sequences of their 16S rRNA gene differ by 9.1–9.7% from the 16S rRNA of *Holospora* species. These symbionts do not induce formation of the connecting piece in the nucleus during division of *Paramecium*.

Here we report a new *Holospora*-like intranuclear bacterium in the macronucleus of ciliate *Paramecium putrinum* originating from Yakutia (Sakha Republic), Russia. Our microscopical observations, phylogenetic analysis based on the 16S rRNA genes, and fluorescence in situ hybridization assays allow to suggest its inclusion as a novel member of genus *Gortzia*. We suggest this bacterium to be classified as a new species *Сandidatus* Gortzia yakutica sp. n.

## Materials & methods

### Sampling & identification of Paramecium

The ciliate *Paramecium putrinum* was originally isolated from a freshwater pond in Yakutia (62°02′N 129°44′E), Sakha Republic, Russia in the summer of 2013. Monoclonal cultures of this species were maintained under standard conditions as described in Sonneborn at the room temperature in lettuce medium inoculated with the bacterium *Enterobacter aerogenes* as food resource (Sonneborn, 1970). The host was identified by the cell morphology, structure of micronucleus and contractile vacuole (Wichterman 1986; Fokin 2010). Living observations and images were made at the Research park of St. Petersburg State University Center for Culturing Collection of Microorganisms with a Leica DM2500 microscope equipped with differential interference contrast (DIC).

The syngens of different *P. putrinum* clones were determined by series of crossing with *P. putrinum* test-clones from two syngens (reproductively isolated groups). All cultures were fed the day before the experiment. Approximately 100 cells of the testing clones were mixed with the equal number of test-clones’ cells. The syngen distribution of tested clones is in good accordance with two clades identified via phylogenetic analysis of the cytochrome oxidase I gene (Tarcz et al. 2014).

### Phenotypical characterization of the symbionts

The infectious capability of symbionts was proved by adding IFs of the bacteria to a non-infected *P. putrinum* culture. Cross-infection experiments were performed with different *P. putrinum* clones from both syngens. *Paramecium* cells containing IFs of the symbiont were concentrated at 4500g for 10 min and homogenized using 1% solution of detergent Nonidet P-40 (Sigma-Aldrich Cat No. 21-3277 SAJ). A small amount of the homogenate was checked at 200x magnification to verify that all ciliate cells had been broken. Equal amounts of homogenate were mixed with recipient *Paramecium* cultures and incubated at the room temperature. Cells were observed at 24 and 48 h post-infection. Additional checks of the mixed cultures were performed four times during the following two months.

### Purification and sequencing of symbionts

The cell culture of *P. putrinum* containing IFs of the symbiont was concentrated and homogenized as stated above. The infectious forms of the endosymbiont were isolated from the homogenate by centrifugation in Percoll density gradient (Sigma-Aldrich Cat No. P1644) as described previously (Rautian and Wackerow-Kouzova 2013). DNA from the purified IFs was isolated using the DNeasy Blood and Tissue kit (QIAGEN Cat No. 69504) using a modified protocol as described previously (Garushyants et al. 2018).

Bacterial universal primers 27F1 (5’-AGAGTTTGATCCTGGCTCAG-3’) and 1492R (5’-GGTTACCTTGTTACGACTT-3’) were used for the amplification of 16S rDNA. PCR products were gel purified and cloned with the TIAN Quick Midi Purification Kit (Tiangen, Beijing, China) following the manufacturer’s recommendations. Purified rDNA inserted in the PTZ57 R/T plasmid vector (InsTAclone PCR Clone Kit, Fermentas, Canada), the recombinant plasmids were transformed to competent cells Trans5α (TransGen Biotech, BeiJing, China). The positive clones were digested with *Hha*I (Fermentas, Thermo Scientific). Clones determined to be unique by the RFLP analysis were sequenced by an automated ABI DNA sequencer (model 373, PE Applied Biosystems).

### Fluorescence in Situ Hybridization (FISH)

Fluorescence in situ hybridization (FISH) with rRNA-targeted probes was performed to visualize the localization of the symbiont. The probe was designed specifically for the new HLB *–* Gyak567 (5’-AGGTAGCCACCTACACA-3’). The sequence was labeled with the cyanine 5 (Cy5) fluorescent dye at the 5’ end. We also used the ALF1b probe for *Alphaproteobacteria* labeled with cyanine 3 (Cy3) as a positive control. *P. putrinum* cell culture containing symbionts was concentrated using centrifugation at 3000 g for 10 min. Cells were fixed in 4% paraformaldehyde in the 1X PBS buffer at 4°C for 3 h shaken every 30 min. The cells were pelleted by centrifugation and washed twice with the PBS solution to remove the residual fixative. The hybridization buffer (0.9 M NaCl, 20 mM Tris-HCI pH 7.2, 0.01 % SDS) and the probes to the final concentration of 5 ng/μl were added. The hybridization was followed by three 20 min post-hybridization washes at 48°C in the washing buffer (0.9 M NaCl, 20 mM Tris-HCI pH 7.2, 0.01 % SDS). All experiments included a negative control without probes to test for autofluorescence. The slides were observed with Leica TCS SP5 confocal laser scanning microscope in The Chromas Core Facility at Saint Petersburg State University.

### Phylogenetic analysis

Sequences of 16S rRNA genes were obtained from GenBank (Benson et al. 2012) for 33 *Rickettsiales*, *Holosporales*, and other related bacteria (see Fig. 5 for the accession numbers). Only sequences longer 1200 bp and shorter than 1500 bp were used. The initial multiple alignment was constructed using mafft v7.407 (REF) with the default settings and then filtered using BMGE v1.12 (Criscuolo and Gribaldo 2010) based on the sequence divergence and gap percentage, yielding reduction from 2040 to 1300 positions. The resulting alignment was additionally trimmed to account for the shortest sequences; to this end, 102 bp from at the 5’-end and 87 at the 3’-end were trimmed, resulting in the final multiple alignment of 1111 bp. RAxML v8.2.12 (Stamatakis 2014) was run on this alignment. A rooted phylogenetic tree was generated using GTRGAMMA model with 1000 bootstraps. The resulting rooted phylogenetic tree was visualized using Interactive Tree of Life v4 (Letunic and Bork 2019). The 16S rDNA sequence of the new HLB, strain YA111-52, was deposited in the GenBank database under the accession number MK209687.1. Sequence similarity was assessed by reciprocal megablast alignment of full GenBank 16S rDNA nucleotide sequences and visualized using ggplot2 and R.

## Results

### Bacterial morphology and localization

Symbiotic bacteria inhabit macronucleus of *P. putrinum* (Fig. 1 A). The infection was stable in several clones of *P. putrinum* for at least three years under laboratory conditions. Uncharacteristically for HLB, a small number of infectious forms could be found in the cytoplasm of the host cell (Fig. 2), suggesting that there is an intermediate state before the symbiont release into the environment. The symbionts were observed in two morphological forms of their life cycle: (1) small (1-2 × 2-4 μm) bacteria undergoing constant binary fissions (reproductive forms, RFs), and (2) long (1-2 × 7-12 μm) infectious forms (IFs). Most observed IFs had straight rod-like shape with tapered ends, and some were slightly curved (Fig. 1 B). The symbionts were never observed in the micronucleus both in stably infected cultures and during the infection process.

**Figure 1.**
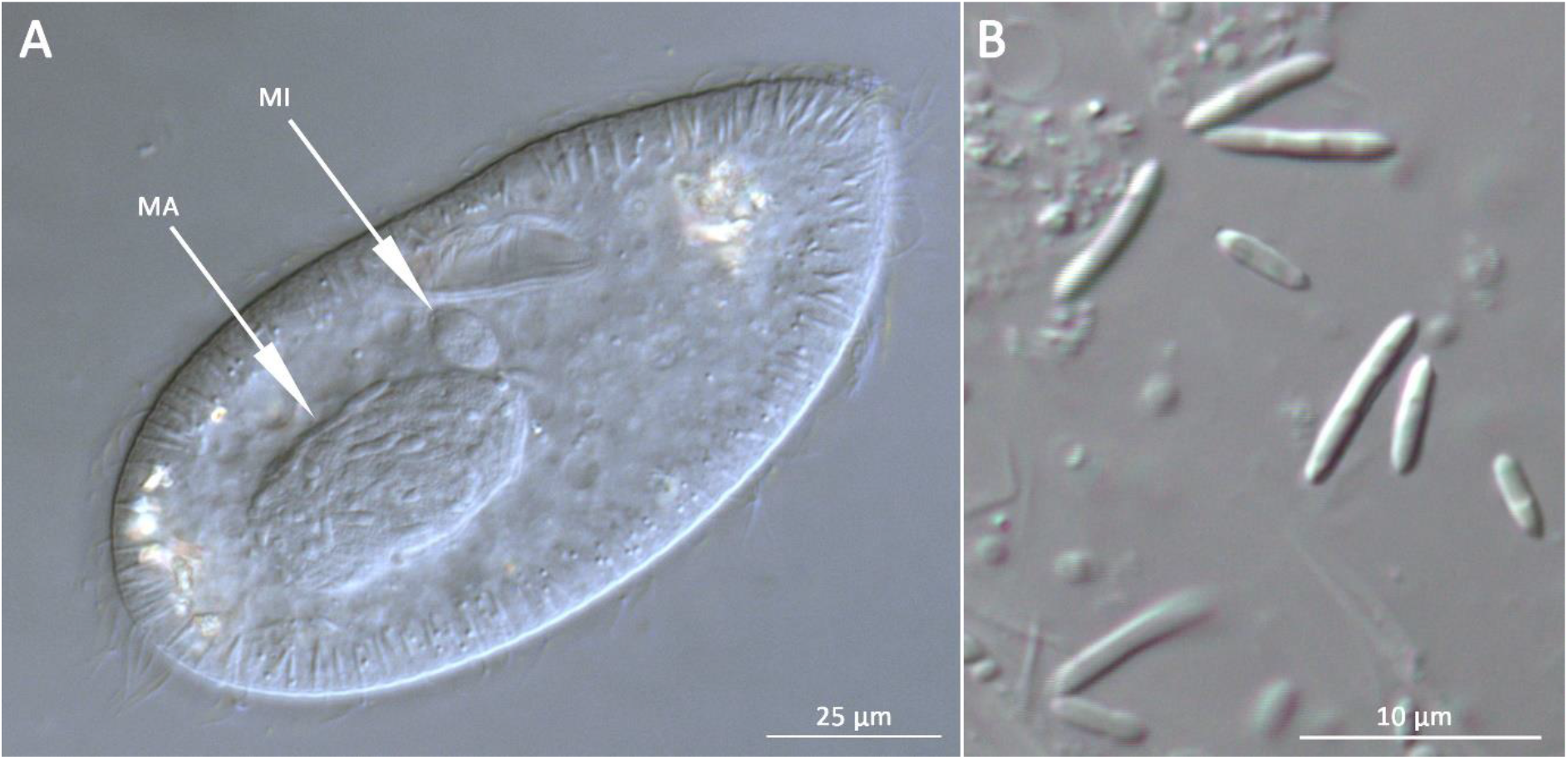
(A) *P. putrinum* with bacteria in the macronucleus; MA – macronucleus, MI – micronucleus; (B) Infectious forms of the new HLB released from the macronucleus.

**Figure 2.**
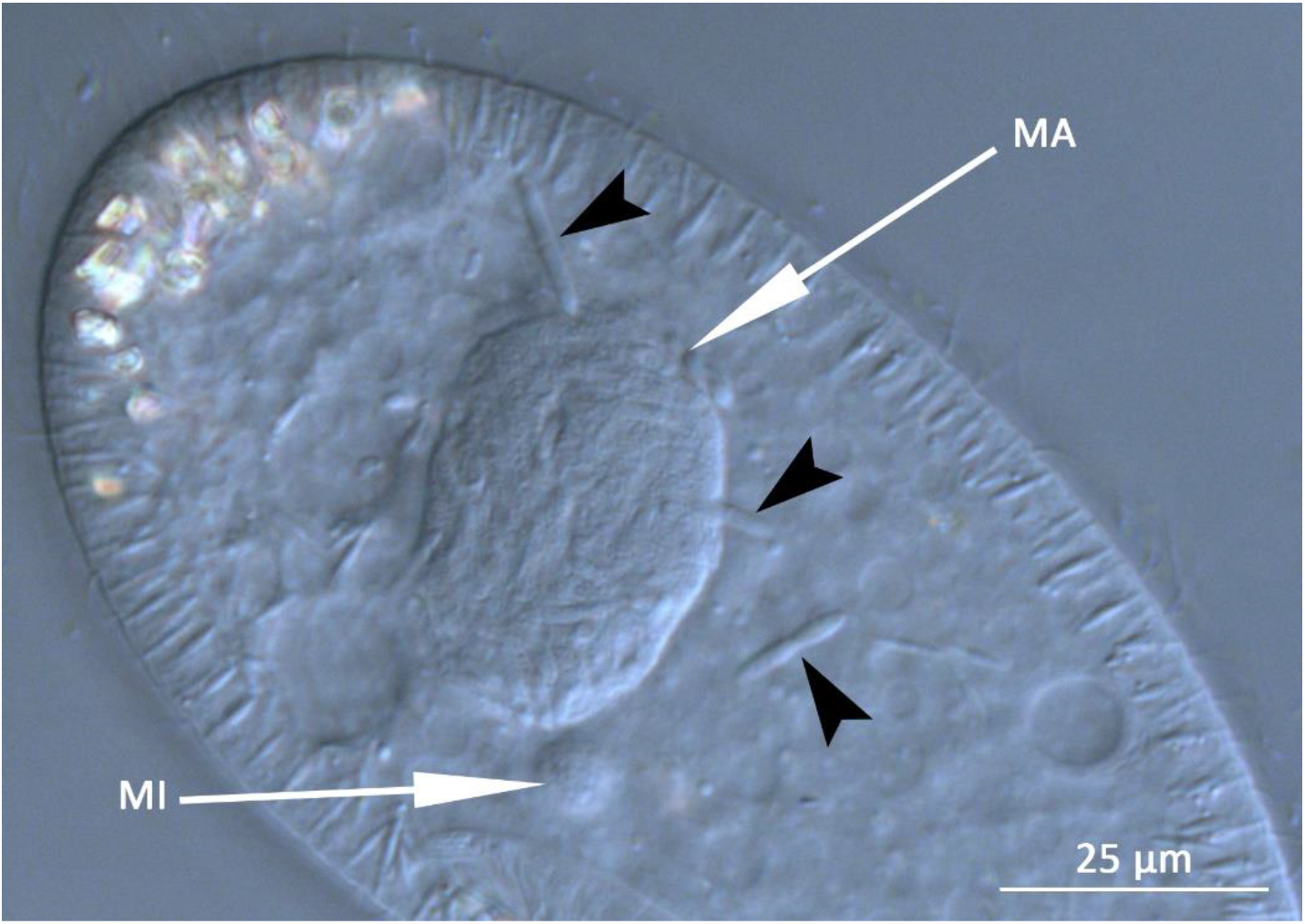
*P. putrinum* with bacteria in the macronucleus and individual infectious forms in the cytoplasm (shown with the black arrowheads). MA – macronucleus, MI – micronucleus.

We never observed formation of the connecting piece during the host cell division (Fig. 3), similar to previously described species from the genus *Candidatus* Gortzia. This feature and the nuclear- and host-specificity also places this new symbiont close to «*H. bacillata*», «*H. curvata*» and *H. sp.* from the macronucleus of *P. putrinum* (Fokin and Gortz 2009).

**Figure 3.**
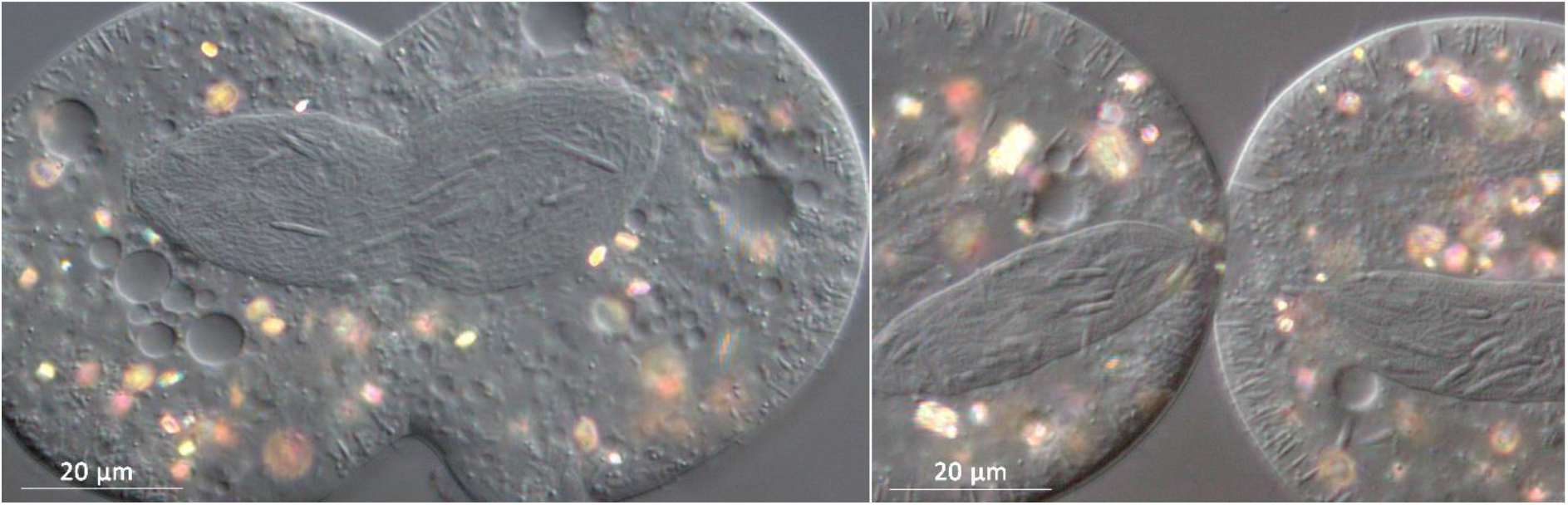
*P. putrinum* during the division process. No connecting piece is observed.

The symbionts are capable of infection aposymbiotic cells of *P. putrinum*. The IFs reach macronucleus and begin to divide in 20-30 hours after infection forming chains of cells characteristic for HLB. Aposymbiotic cells of two *P. putrinum* clones belonging to different syngens were experimentally infected by IFs of the new symbiont. Infection of both clones remained stable for at least two months.

### Molecular characterization

A 1393 bp long 16S rDNA sequence was obtained from the new HLB and deposited at GenBank under the accession number MK209687.1. Using the FISH technique with the Gyak567 probe we detected a large number of bacteria in the macronuclei of *P. putrinum* (Fig. 4B). The sequence-specific probe Gyak567 bound to bacteria inside the macronucleus in our FISH experiments, thus demonstrating that the characterized 16S rRNA gene sequence derives from the HLB. The ALF1b probe was used as a positive control (Fig. 4A).

**Figure 4.**
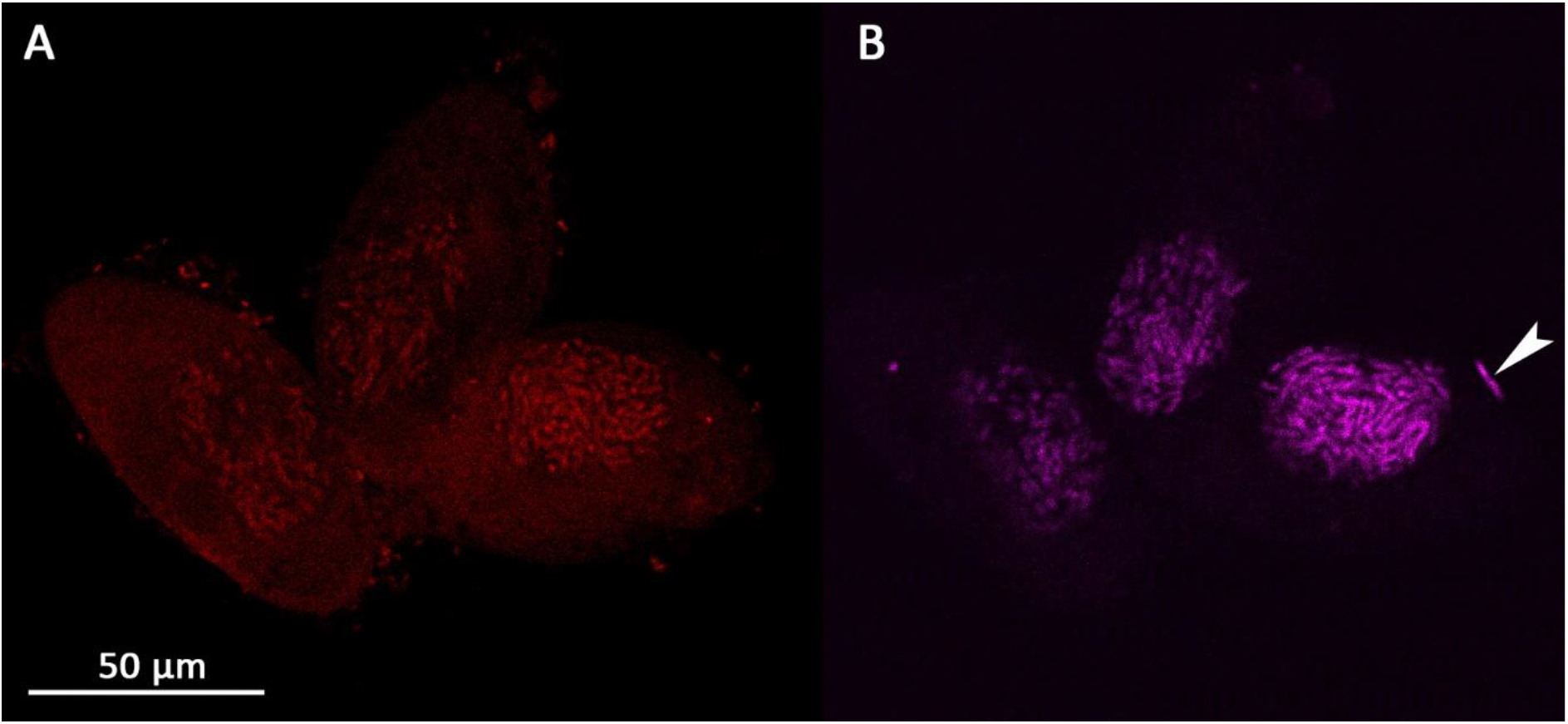
Cells of *P. putrinum* with symbionts in macronuclei labelled with the probes ALF1b (A) and Gyak567 (B). Single IF lying outside the macronucleus is shown with the white arrowhead.

### Phylogenetic Analysis

The phylogenetic analysis confidently places the new HLB within the *Gortzia* branch as a sister taxon to two other *Gortzia* species, *Candidatus* G. infectiva and *Candidatus* G. shahrazadis, macronuclear symbionts of *P. jenningsi* and *P. multimicronucleatum*. However, the level of sequence divergence of 3% and 3.5% of the new HLB with *Candidatus* G. shahrazadis and *Candidatus* G. infectiva respectively suggests that the new HLB is a separate species within the HLB clade and the genus *Candidatus* Gortzia. Two previously described *Candidatus* Gortzia species have 1.08% of divergence in their published 16S rDNA sequences. The difference of the new HLB with species of *Holospora* genus varies between 9% and 11% (Fig. 6). Since HLBs are obligate symbionts and are not cultivable outside host cells, a complete culture-dependent characterization cannot be provided; hence, we propose the provisional name “*Candidatus* Gortzia yakutica”.

**Figure 5.**
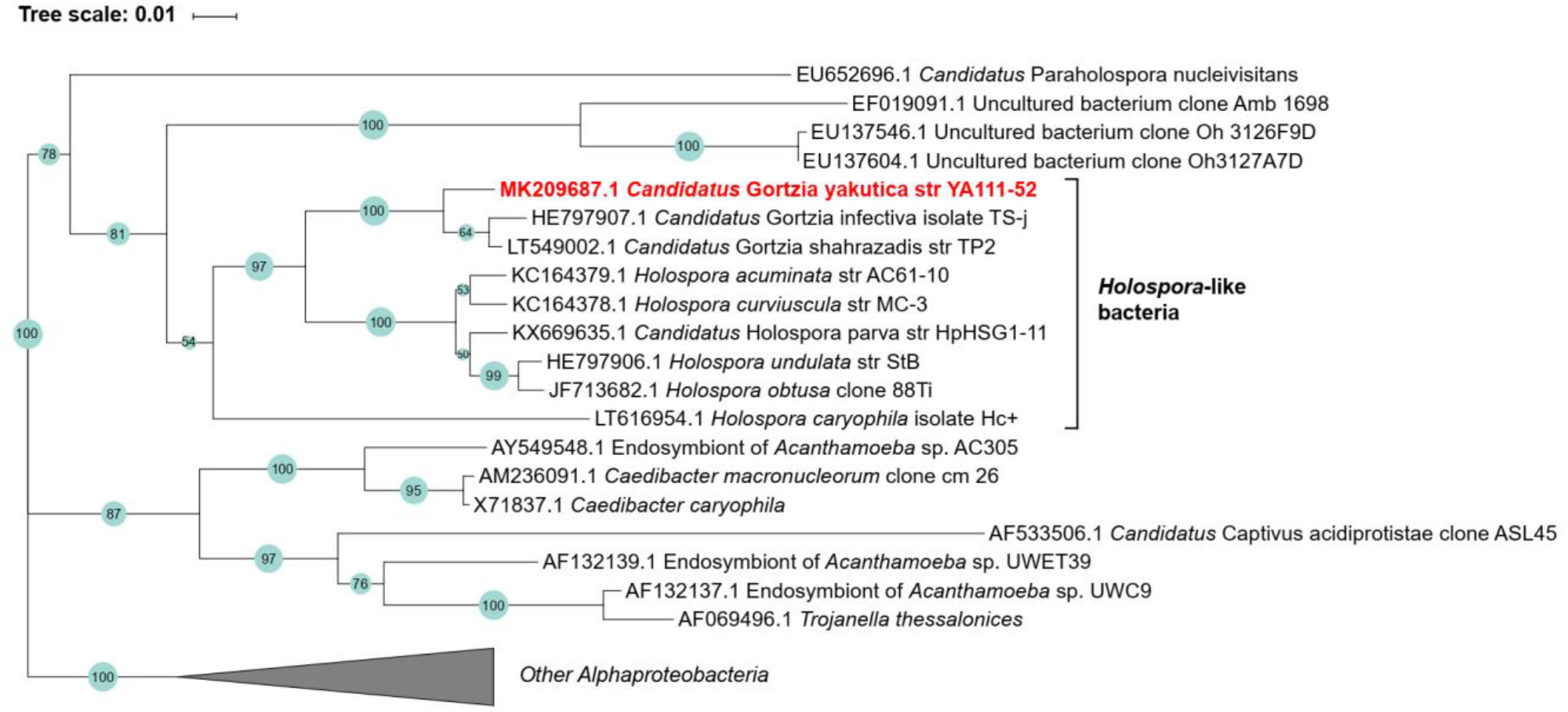
Maximum likelihood phylogenetic tree of the order *Holosporales*. Bootstrap support values are shown on each branch.

**Figure 6.**
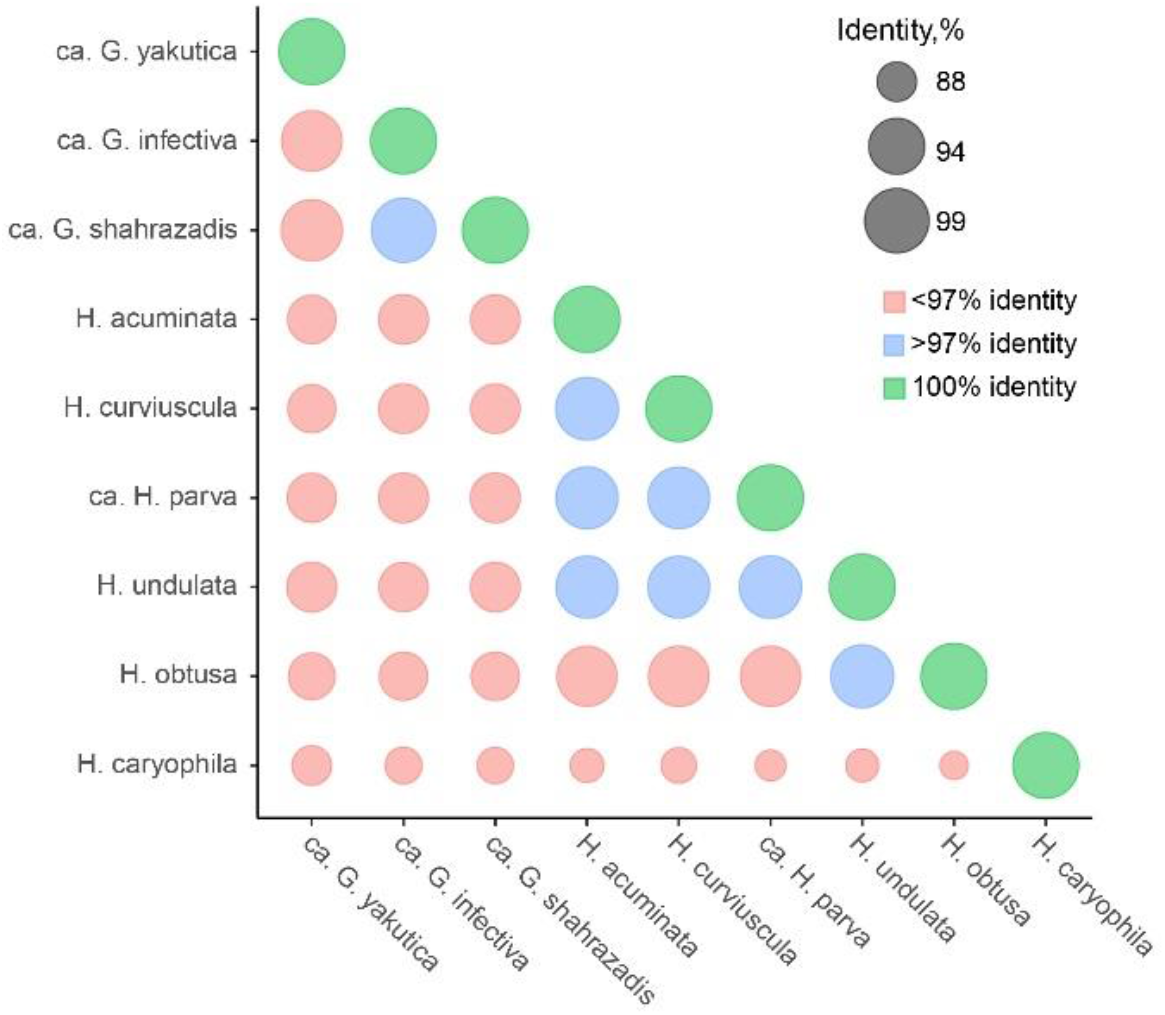
Divergence of *Holospora*-like bacteria based on 16S rDNA gene. Nucleotide blast identities of full 16S sequences are used as a measure of sequence similarity.

The phylogenetic tree shows a convincing monophyly of all HLB including genera *Holospora* and *Candidatus* Gortzia (Fig. 5). The sister clade to HLB includes other symbionts, e. g. genus *Caedibacter.*

## Discussion

Here, we report a new *Holospora*-like bacterium from the macronucleus of *P. putrinum*. All features of this bacterium, such as morphology, intracellular localization, complex life cycle, host and nuclear specificity, and infectivity, indicate a close relation of this symbiont to other *Holospora-*like bacteria. It is possible that this species was described previously by Fokin (S. Fokin and Gortz 2009) as “*Holospora* sp. from macronucleus of *P. putrinum*”, but that culture had been lost precluding a detailed characterization.

The phylogenetic analysis based on the 16S rRNA gene sequence also shows that the symbiont is close to other *Holospora*. Recently described macronuclear symbionts from *P. multimicronucleatum*, *Candidatus* G. infectiva and *Candidatus* G. shahrazadis are the closest relatives of the new symbiont and together they form a sister clade to other *Holospora* species. Similar to *Candidatus* G. infectiva and *Candidatus* G. shahrazadis, the new symbiont does not induce formation of the connecting piece during the host division. Consequently, we assign this symbiotic bacterium to the genus *Gortzia* and name it *Candidatus* Gortzia yakutica sp. nov.

Notwithstanding that our study supports the *Gortzia* clade, the issue of creating a new genus is worth discussing. The HLB clade is clearly monophyletic, and all HLB appear to be very similar in their morphological and physiological features. The analysis of 16S rRNA gene sequences is widely used to make decisions on bacterial taxonomy, despite obvious limitations caused by frequent multiplicity of 16S rRNA genes in a genome resulting, in some cases, substantial variation in the 16S rRNA sequences within the same genome, as well as varying similarity level within a species, a genus, or a family. An approximate threshold of 97% 16S rRNA similarity was previously reported to delineate a bacterial species (Větrovský and Baldrian 2013). For many phenotypically characterized HLB the complete 16S gene has not been sequenced yet. It has been recently demonstrated that complete genome sequences could be used to clearly define bacterial species, effectively settling a long-standing theoretical debate (Jain et al. 2018; Moldovan and Gelfand 2018). Thus, we can conclude that we would be in a much better position to infer the phylogenetic relationships of the *Gortzia* clade when complete genomes of *Gortzia* spp. become available.

## Description of Gortzia yakutica sp. nov

Gortzia yakutica (Gor’tzi.a ya.ku’ti.ca; N.L. fem. n. Gortzia, in honour of Professor emeritus Hans-Dieter Görtz; N.L. fem. adj. *yakutica*, of or belonging to Republic of Yakutia, the name of the region where the bacterium was first collected).

Obligate macronuclear endosymbionts of the free-living ciliate *P. putrinum*, occasionally can be found in the cytoplasm. Sampled from the fresh-water pond in Republic of Yakutia, Russia. Has two life stages: small reproductive forms (1-2 by 2-4 μm) and long infectious forms (1-2 by 7-12 μm, rod-shaped with tapered ends, sometimes slightly curved). No formation of the connecting piece was observed. Basis of assignment: SSU rRNA gene sequence (GenBank accession number: MK209687.1) and positive match with the species-specific FISH oligonucleotide probe Gyak567 (5’-AGGTAGCCACCTACACA-3’). Unculturable outside of host cells.

